# Real-time fine aerosol exposures in taconite mining operations

**DOI:** 10.1101/263442

**Authors:** Tran Huynh, Gurumurthy Ramachandran, Harrison Quick, Jooyeon Hwang, Peter C Raynor, Bruce H Alexander, Jeff H Mandel

## Abstract

Respiratory health effects such as mesothelioma, silicosis, and lung cancer have been shown to be associated with working in the taconite mining industry. Taconite workers may also have elevated risks from cardiovascular disease (CVD), although the relationship of CVD to dust exposures at these mines has not been well-studied. Motivated by evidence from environmental epidemiological studies and occupational cohorts that have implicated the effects of fine particulates with increased risk of cardiovascular diseases, we conducted an air monitoring campaign to characterize fine aerosol concentrations at 91 locations across six taconite mines using an array of direct-reading instruments to obtain measurements of mass concentrations (PM2.5 or particles with aerodynamic diameter less than 2.5 μm, and respirable particulate matter or RPM), alveolar-deposited surface area concentrations (ADSA), particle number concentrations (PN), and particle size distributions. To analyze these data, we fit a Bayesian hierarchical model with an AR(1) correlation structure to estimate exposure while accounting for temporal correlation. The highest estimated geometric means (GMs) were observed in the pelletizing and concentrating departments (pelletizing maintenance, balling drum operator, and concentrator operator) for PM2.5 and RPM. ADSA and PN generally had highest GMs in the pelletizing department, which processed large amounts of powder-like particles into iron pellets. The within-location variability (GSD_WL) generally ranged from 1 to 3 for all exposure metrics, except for a few locations which indicated changes of activities that caused the exposures to change. Between-location variability (GSD_BL) estimates were generally higher than GSD_WL, indicating larger differences in exposure levels at different locations between mines than at individual locations over the course of several hours. Ranking between PM2.5 and RPM generally agree with each other, whereas ADSA and PN were more consistent with each other, with some overlap with PM_2.5_ and RPM. Differences in ranking these groups may have potential implication for occupational epidemiological studies that rely on exposure information to detect an exposure-response relationship for various job groups. Future epidemiological studies investigating fine aerosol exposures and health risks in occupational settings are encouraged to use multiple metrics to see how they influence health outcomes risk.

## INTRODUCTION

Mining on the Iron Range in northeastern Minnesota has been ongoing since the late 1800s, when pockets of high-quality iron ore were discovered. By the end of World War II, much of the natural ores were depleted and mining of low-grade ores, called taconite, on the 130-mile Mesabi Iron Range began in the 1950s (MDNR, 2014). Taconite is a hard, dense, sedimentary rock that contains 20 to 30% iron content (MDNR, 2014). Unlike the natural ores, which can be directly used in steel mills, taconite rocks require extensive processing in order to obtain the iron concentrate. Taconite mining and processing releases mineral fibers and particles that have been a public concern since the 1970s (Wilson et al, 2008; Axten and Forster, 2008; Berndt and Brice, 2008). Early health studies did not find increased risk of mortality associated with occupational dust exposure (Clark et al., 1980; Higgins et al., 1983; Cooper et al., 1988; 1992), nor risk of lung cancer and mesothelioma from amphibole cleavage fragments exposure (Gamble and Gibbs, 2008). More recent studies, however, reported elevated rates of mesothelioma in Minnesota counties near taconite processing facilities (MDH, 1999, 2003; Case et al., 2011). Concerns about the potential excess rates of mesothelioma in this cohort led the Minnesota State Legislature in 2008 to fund the Taconite Workers Health Study (TWHS) to evaluate mesothelioma, lung cancer, and non-malignant respiratory disease (University of Minnesota, 2014). Since then, studies on exposure assessment (Hwang et al., 2013 and 2014), mortality (Allen et al, 2014; Mandel et al.. 2016), cancer incidence (Allen et al.. 2015), lung cancer case-control (Allen et al., 2015), mesothelioma case-control (Lambert et al.. 2015) and pulmonary function tests (Odo et al. 2013) have been reported. These studies report that taconite workers are more likely to have mesothelioma, lung, laryngeal, stomach, and bladder cancer and cardiovasculare disease (specifically, hypertensive heart disease and ischemic heart disease) than the general population. Also, length of employment in the taconite industry and potentially exposures to generated elongate mineral particles (EMP) were associated with risk of mesothelioma.

While much of the public concern has surrounded the increased risk of respiratory diseases such as silicosis and mesothelioma, little attention has been paid to non-respiratory adverse health outcomes resulted from working in dusty taconite mining operations. Increased risk of cardiovascular diseases from short-term and long-term exposures to environmental air pollution, particularly fine particulate matters (PM) and ultrafine particles (UFP), are well documented (e.g., Schwartz, 1990; Dockery et al., 1993 and 2005; Pope et al., 2004; EPA, 2014). Similar findings were also reported in occupational cohorts, including workers in aluminum smelters (Costella et al., 2014, Neophytou, 2014 and 2016), construction (Toren et al., 2007), highway maintenance (Meier et al. 2014), and the textile industry (Gallenher, 2012).

Epidemiological studies investigating the association of air pollution, particularly fine and ultrafine particles – and adverse health effects typically use mass and/or particle number concentration for their exposure metrics (e.g, Meier et al, 2014, Dockery et al, 1994). Evidence of the toxicity of UFP suggests that surface area may be more closely related to health effects (Oberdorster, 2005; Maynard and Maynard, 2003; Maynard and Aitken, 2007), yet very few health studies have explored alternative exposure metrics such as surface area concentration (Meier et al., 2014). Of the occupational exposure studies examining the relationship between mass, particle number (PN), and alveolar-deposited surface area (ADSA) concentrations, rankings of exposure groups were shown to differ by metrics where groups with high mass concentration may rank lower in the PN and SA metrics (e.g., Ramachandran et al, 2005; Park et al., 2010, Heitbrink et al., 2009). Therefore, simultaneous measurements of multiple metrics is often recommended for assessing fine and ultrafine particle (including nanoparticle) exposures (Park et al. 2010, Baldauf et al., 2015).

The goal of this study is to characterize fine aerosol exposures using multiple metrics in taconite mining operations using direct-reading instruments. We begin by reporting the estimated area concentrations at various locations where similarly exposure groups typically work within the mines. We then shift our focus to examining the relationship between exposure metrics in taconite mining settings to better understand how it might influence the ranking of exposure groups. Finally, we offer concluding remarks regarding implications for future occupational epidemiological studies.

## METHODS

### Taconite mining process

The goal of the taconite mining process is to crush the rocks into small granules so that the iron-bearing particles can be separated from the rest. The process comprises four major steps: extracting, crushing, concentrating, and pelletizing. Taconite boulders are unearthed by drilling and blasting the ground with explosives and hauled to a crusher building where they are crushed into small rocks in ∼10 cm in diameter by the primary crushers and a series of fine crushers. In the concentrating department, conveyors move the crushed rocks underneath the stockpile to the rod mills where the ore is mixed with water and ground to the size of sand particles. The slurry discharged from the rod mills is pumped through the magnetic separators that separate the iron-bearing materials from non-iron particles called ‘tailing’. The magnetic ore slurry is pumped to the ball mills that further reduce the ores to a fine powder (0.1-30 µm). A series of magnetic separators called roughers and finishers further concentrate the powder-like materials. The flotation processes use chemicals, air, and mechanical agitation to remove the fine silica particles from the settling iron-bearing particles. In the pelletizing department, disc filters are used remove most of the water from the concentrates by vacuum pumps. The filter cake is mixed with a binding agent (bentonite and/or limestone) that helps to hold the pellets together during baking. The mixture is then fed into rotating discs called balling drums which roll the mixture into marble-size pellets in ∼1 cm in diameter. The pellets are transferred into a kiln where they are heat-hardened and cooled. The finished pellets are loaded onto trains at loading pockets for shipment (EPA, 2003; Minorca Mine Booklet).

#### Sampling strategy

Real-time area monitoring of aerosols was conducted from January 2010 to May 2011 at six mines located along the Mesabi Iron Range in northeastern Minnesota in conjunction with personal particulate exposure sampling conducted by Hwang et al (2013). Samples were taken at work locations that corresponded to the similar exposure groups (SEGs) established for the personal exposure assessment to taconite dust components. Hwang et al. (2014) assembled almost 200 job titles from the Mine Safety and Health Administration (MSHA) Mine Data Retrieval System, jobs from a previous study by Sheehy (1986), industrial hygiene and human resources databases of the three companies currently operating the mines, and *Job Descriptions and Classifications* published by the Reserve Mining Company (1974). These job titles were then condensed to 28 SEGs. For the real-time area monitoring we used 16 SEGs because we did not sample the basin/mining operators who primarily work outside. We also consolidated mobile maintenance-related jobs (e.g., boiler technician, carpenter, electrician, lubricate technician, plumber, repairman) into one maintenance technician SEG because some of these workers are highly mobile and the maintenance or carpenter shop is where these people may consider their home base. Table 1 provides brief descriptions of the tasks and number of locations for each SEG. A total of 92 locations were sampled across six mines.

**TABLE 1:**
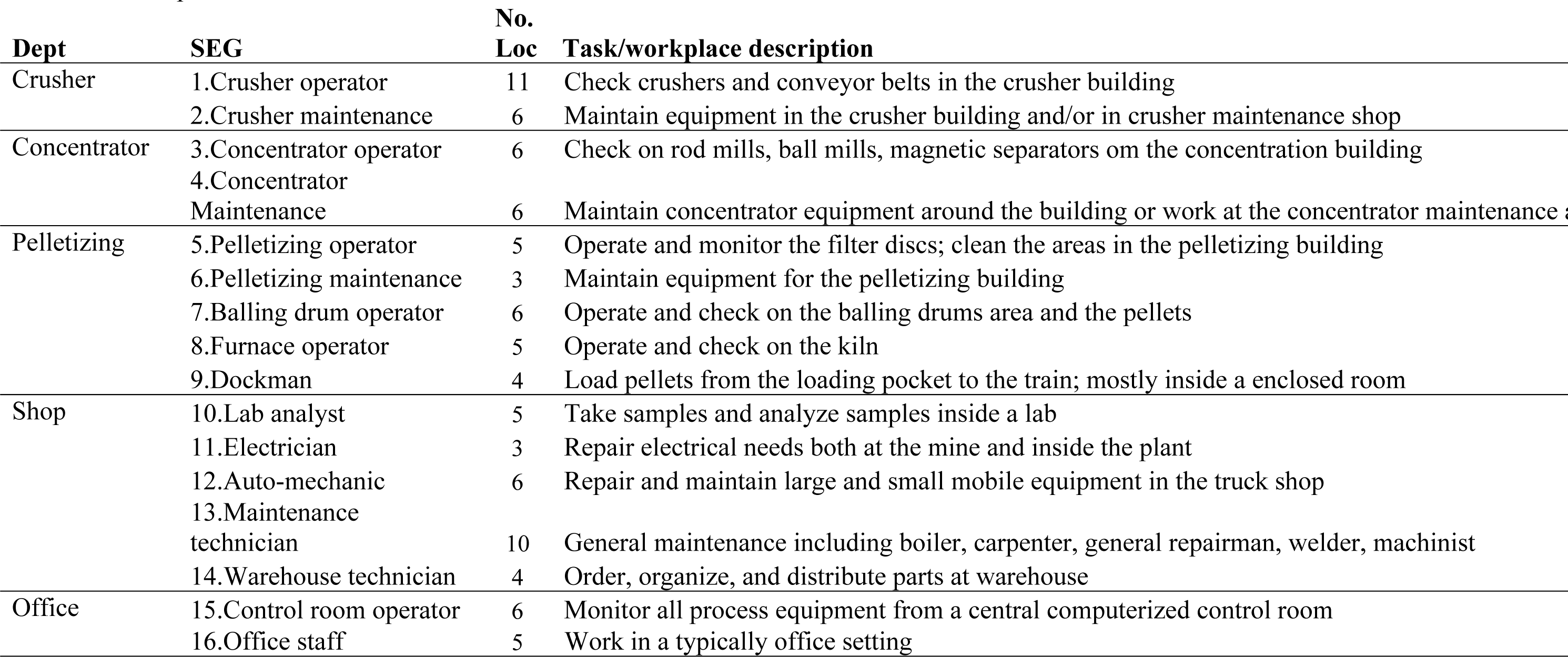
Description of the tasks and locations where area aerosol concentrations were measured.

Area measurements were taken at least once per location per SEG per mine. Some SEGs that covered a large workspace (e.g., the crusher building) were sampled at multiple locations. At each sampling location, a respirable gravimetric pump (Apex Pro pump, Casella Inc., Amherst, NH, USA) and five direct-reading instruments measuring mass, surface area, particle number concentration, and particle size distribution were run simultaneously for approximately 4 hours with 1 minute averaging time (for the direct-reading instruments). Gravimetric samples were sent to an accredited laboratory for analysis using NIOSH 0600 Respirable particulates not otherwise regulated gravimetric. Two photometer air monitors (DustTrak, Model 8520, TSI, Inc., MN, USA) with PM_2.5_, and respirable particulate matter (RPM) sampling inlets were used to obtain corresponding size selective mass concentration in mg/m^3^. A condensation particle counter (CPC) (PTrak, Model 8525, TSI, Inc., MN, USA) measured particle number (PN) concentration (expressed in particles/ cm^3^) in the size range from ∼0.02 µm — ~1.0 µm. An optical particle sizer (Aerotrak, Model 9306, TSI, Inc., Shoreview, MN) counted and sized aerosol particles into various size bins (0.3 µm -0.5 µm, 0.5 µm -1.0 µm, 1.0 µm -3.0 µm, 3.0 µm -5.0 µm, 5.0 µm -10 µm, and >10 µm). An additional size bin (0.02 – 0.3 µm) was obtained by subtracting the CPC total particle count from the first and second bin (0.3 µm -0.5 µm, 0.5 µm -1.0 µm) of the optical particle sizer. A surface area monitor (Aerotrak 9000,TSI, Inc., MN, USA) measured surface area of the particles 0.01-1.0 µm that deposit in the alveolar region of the lung (alveolar-deposited surface area or ADSA). Concentration is expressed in (µm^2^/cm^3^).

We collected gravimetric respirable measurements at each location with the intention of calibrating the DustTrak monitors using the procedure described in Ramachandran (2000). Laboratory analysis returned with about 68% of the samples below the detection of limit (LOD). The majority of < LOD results were from relatively clean environment such as the shops and offices and a few locations within the crusher, concentrator and pelletizing departments. The LODs ranged from 0.11 to 0.18 mg/m^3^. The average correction factor (CF) for RPM samples above the LOD was 1.13 with a range from 0.25-3.05. For filter samples below the LOD, we assigned a value of LOD/2 and computed the correction factor. The average CF for samples below the LOD was 9.77with a range from 0.24 to 186. The high variability in the CFs led us to believe that the raw DustTrak measurements would be more reliable than the corrected measurements. While calibration of this DustTrak model with a validated instrument was recommended (Yanosky, 2001), the use of gravimetric RPM measurements LODs for calibration could potentially vastly overestimate the raw DustTrak measurements depending on the locations. This DustTrak model has been shown to provide reasonably precise estimates (R^2^=0.859) for PM_2.5_ (Yanosky, 2001, Chung et al. 2001). We decided to use the raw DustTrak measurements for analysis knowing that it might be slightly biased compared to a reference method.

### Statistical analysis of the data

In order to seamlessly obtain uncertainty estimates in addition to the standard estimates of GM and GSD, a Bayesian hierarchical model was used to estimate exposure while accounting for the temporal correlation present in the data to reduce biased estimates of the standard errors [Klein Entink et al, 2015]. While a classical AR1 regression model accounting for temporal correlation could be used, the advantage of the Bayesian approach is that it can seamlessly provide uncertainty estimates of the GM and GSD and impute any missing values in the data.

In the Bayesian model, we let *Y_ml(g)t_* denote the concentration of metric *m* observed in location *l* of exposure group *g* at time *t*, where *m* = 1, …, 4 (PM_2.5_, RPM, SA, and PN), *l* = 1, …, L_g_. We put the subscript for group, g, in parentheses to signify that location is *nested within* group, location sampled for each exposure group ranged from 3 to 11 (see Table 1), g = 1, …, 16, and t = 1, …, 240. We fit a Bayesian hierarchical model with an AR(1) (autoregressive order 1) correlation structure where

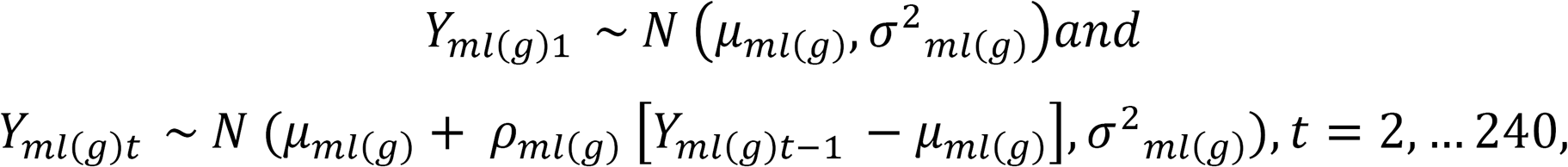

Where *µml(g)* denotes the log GM of metric *m* corresponding to the l^th^ location, *σmi(g)* denote the overall log GM and between-location log GSD of metric *m* corresponding to the *l^th^* location, and *µml(g)* accounts for the temporal correlation in the data (e.g., *µml(g)*= 0 indicates temporal independence and *µml(g)*= 1 indicates perfect correlation between the first and second measurement). To learn about between-location variability, we let *µml(g)*∼*N*, θ_mg_, τ^2^_*mg*_where *θ_mg_* and *τ_mg_* denote the overall log GM and between-location log GSD of metric *m* for exposure group *g,* respectively. The specification of our hierarchical model was then completed by assigning prior distributions to the various model parameters: a non-informative flat prior for *θ_mg_*, vague inverse gamma priors for *τ^2^_mg_* and *σ^2^_ml(g)_* and a uniform(-1,1) prior for *ρ_ml(g)_*. These conventional vague priors were chosen so that they will have little influence over the data. They were also used in the analysis for all four metrics (PM_2.5_, RPM, SA, and PN). For each parameter, we have the posterior distribution of 10,000 iterations. All analyses were carried out using the RJAGS package (Plummer, 2003) in the R statistical software (R Core Team, 2017); for more information on Bayesian methods, see Carlin and Louis (2009).

Prior to fitting the Bayesian model, we plotted the measurements for each metric to see how they tracked each other at each location. We also computed mean at each location and estimated the correlation for between metrics.

## RESULTS

Figure 1 shows an example of how closely the various metrics tracked each other. The correlation plots of average concentrations at each location for different combination of metrics were demonstrated in Figure 2. PM_2.5_ and RPM average concentrations are highly correlated to each other (Pearson’s correlation = 0.98), followed by ADSA and PN correlation (R=0.77). Table 2 reports the estimated geometric means (GM), within location variability (GSD_BL), and between-location variability (GSD_WL) by SEG and metric. The PM_2.5_ concentrations ranged between 0.008 and 0.269 mg/m^3^ and the three highest concentrations (in descending order) were observed in the locations corresponding to the pelletizing maintenance, balling drum operator, and concentrator operator SEGs, respectively. RPM ranged between 0.009 to 0.38 mg/m^3^. The three highest areas were the same as for PM_2.5_ but in a slightly different descending order: locations corresponding to the pelletizing maintenance, concentrator operator, and balling drum operator SEGs. The SA concentration ranged from 3.66 – 312.34 µm^2^/cm^3^. The highest levels were observed in the locations corresponding to the furnace operator, pellet maintenance, and dockman SEGs. The estimated particle numbers ranged from 848 – 111,282 particles/cm^3^ with highest levels observed in the locations corresponding to the pellet maintenance, balling drum operator, and furnace operator SEGs, respectively. There were eight locations that did not have all four instruments (denoted by *); a separate analysis that only included locations with all four metrics (Table 2 in the Supplement) found that exclusion of these locations did not substantially affect our top three ranking, hence our decision to include those eight locations in the analysis. Figure 3 shows that there is high uncertainty around the GM estimates in the highest exposure groups in the pelletizing department and less uncertainty in the low exposure groups.

**TABLE 2:**
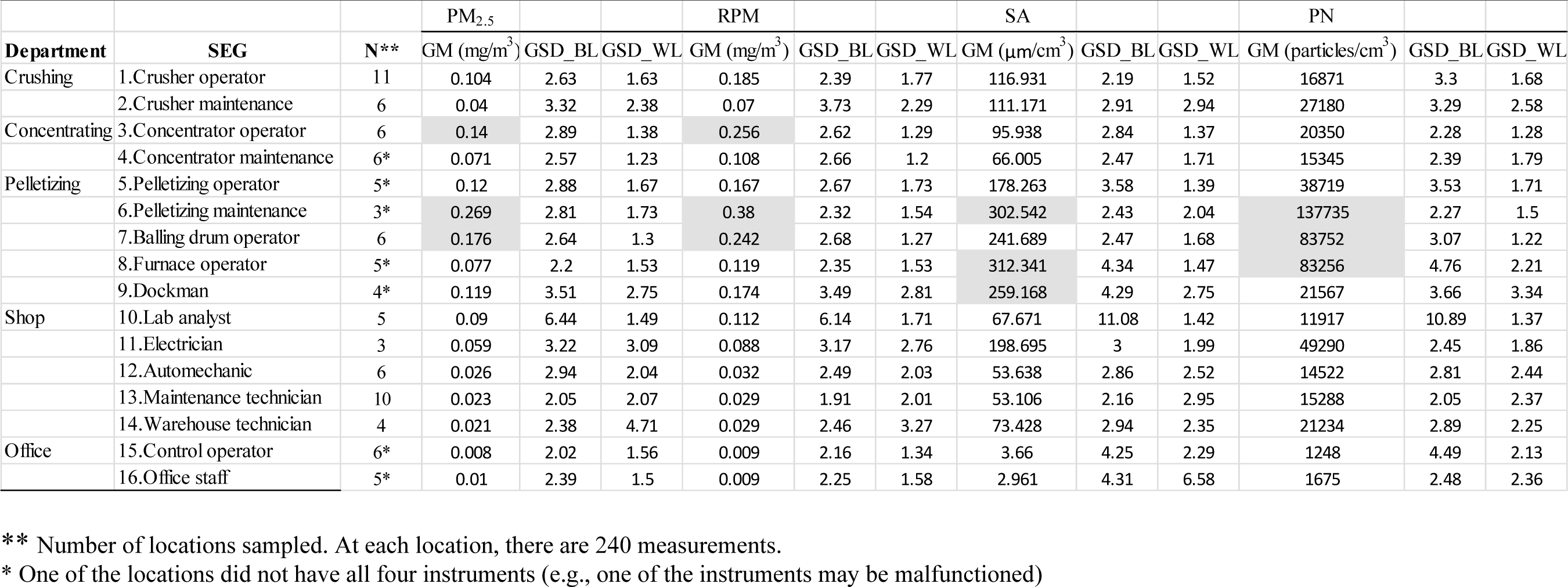
Geometric mean (GM) and geometric standard deviation between locations (GSD_BL) and within locations (GSD_WL) for PM_2.5_, respirable particulate matter (RPM), alveolar-deposited surface area (ADSA), and particular number concentration by exposure groups (SEG). The top three ranking is highlighted in gray.

**Figure 1:**
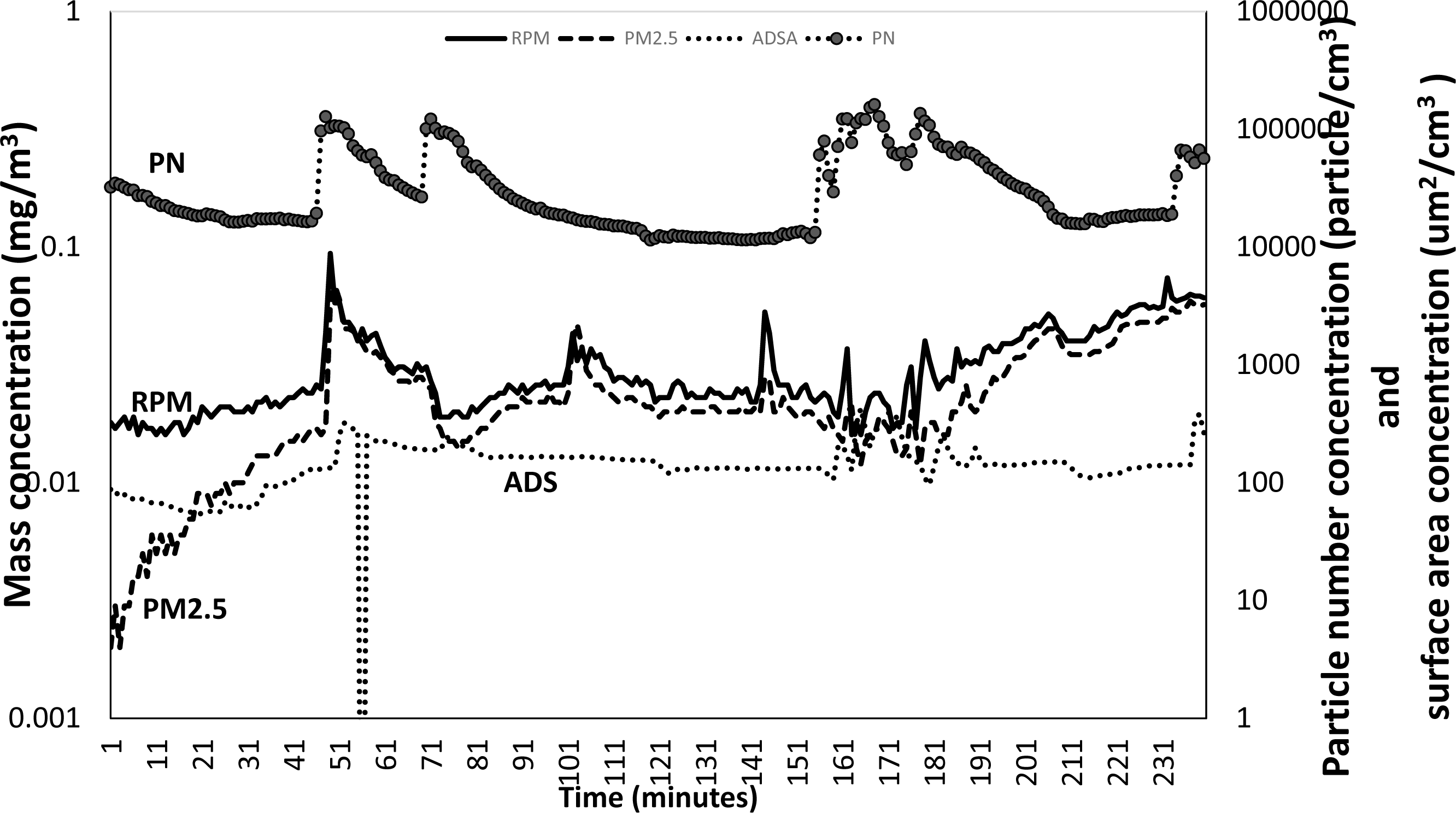
Real-time measurements of PM_2.5_, respirable particulate matter (RPM), alveolar-deposited surface area (ADSA), and particular number concentration (PN) by exposure groups (SEG) at a warehouse.

**Figure 2:**
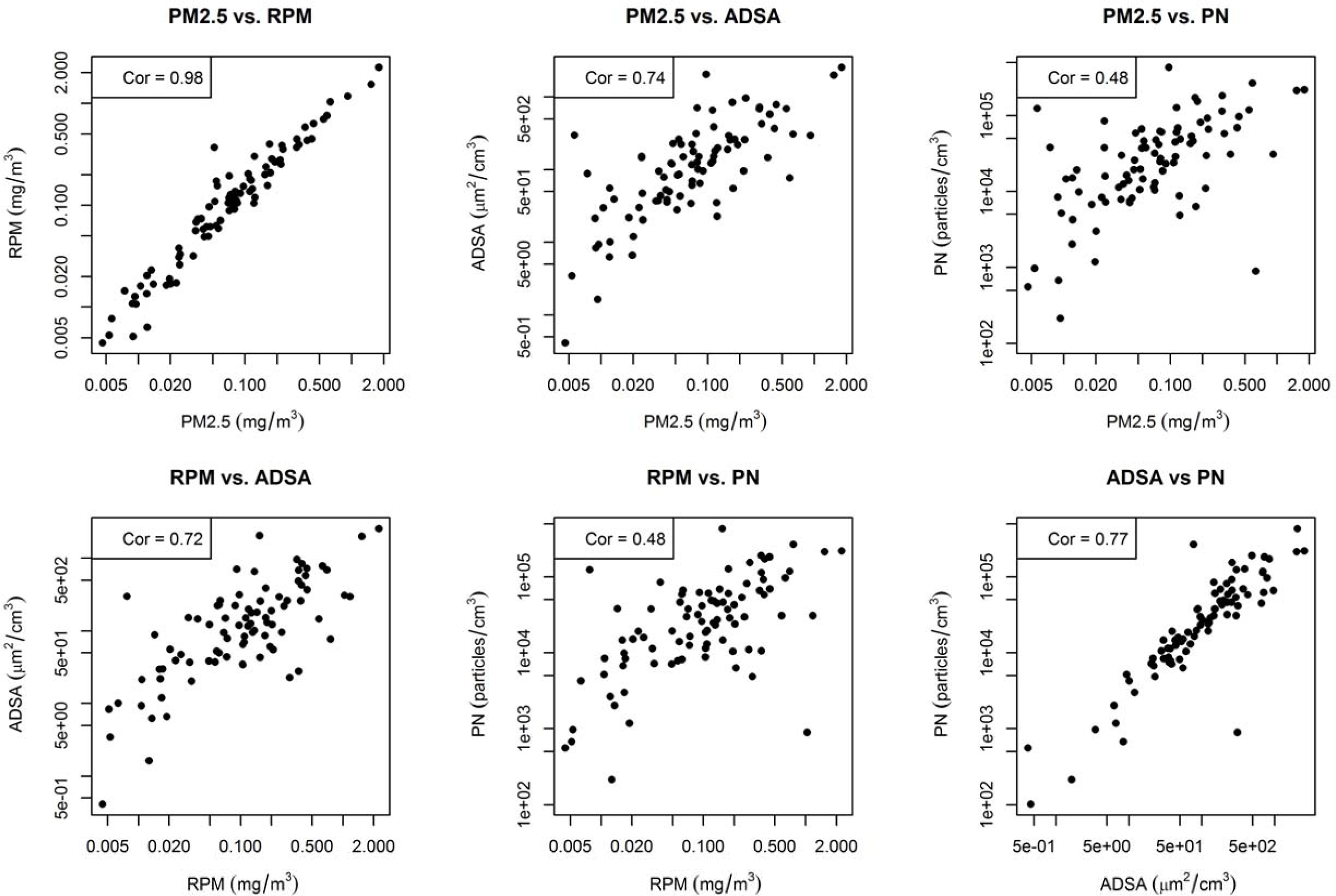
Correlation of average aerosols concentrations at each location for different combination of metrics.

**Figure 3.**
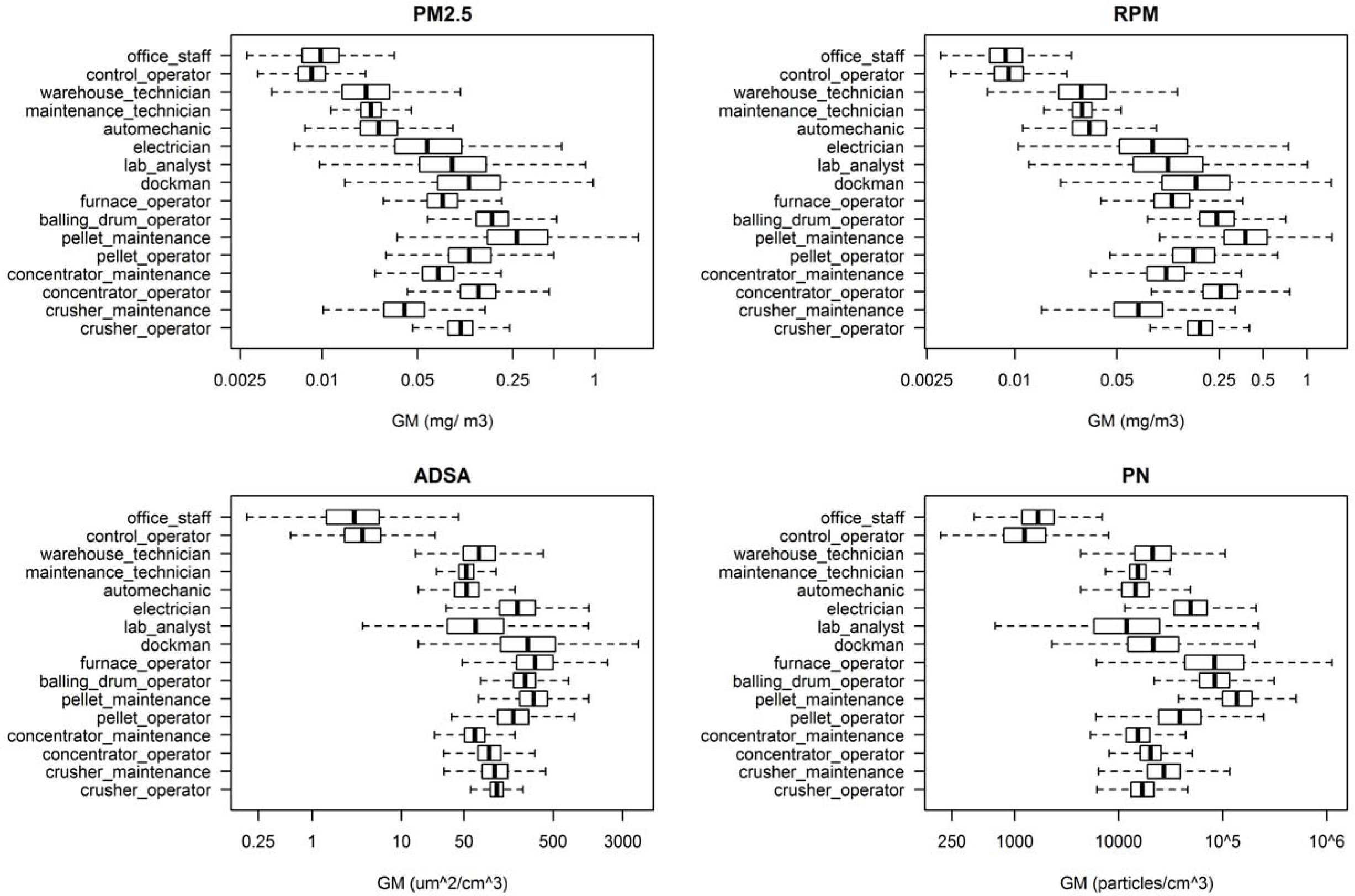
A boxplot showing the level of uncertainty of the geometric mean (GM) estimates by PM_2.5_, respirable particulate matter (RPM), alveolar-deposited surface area (ADSA), and particular number concentration and by exposure groups (SEG).

The GSD_ WL (within location variability) for all locations and metrics were less than 3 except for a few locations such as the warehouse for PM_2.5_ or the office staff for SA, and dockman for PN which also exhibit high level of uncertainty in Figure 4. The GSD_BL (between location variability) in Figure 5 were much higher than GSD_WL in many cases, even as high as 11 for lab analyst SA and PN. It should be noted that the uncertainty intervals for GSD_WL are typically smaller than GSD_BL and, in both cases, the credible intervals get smaller as the GSDs decrease. The GSD_BL and the GSD_WL generally track each other for PM_2.5_ and RPM while those of SA and PN track each other.

**Figure 4:**
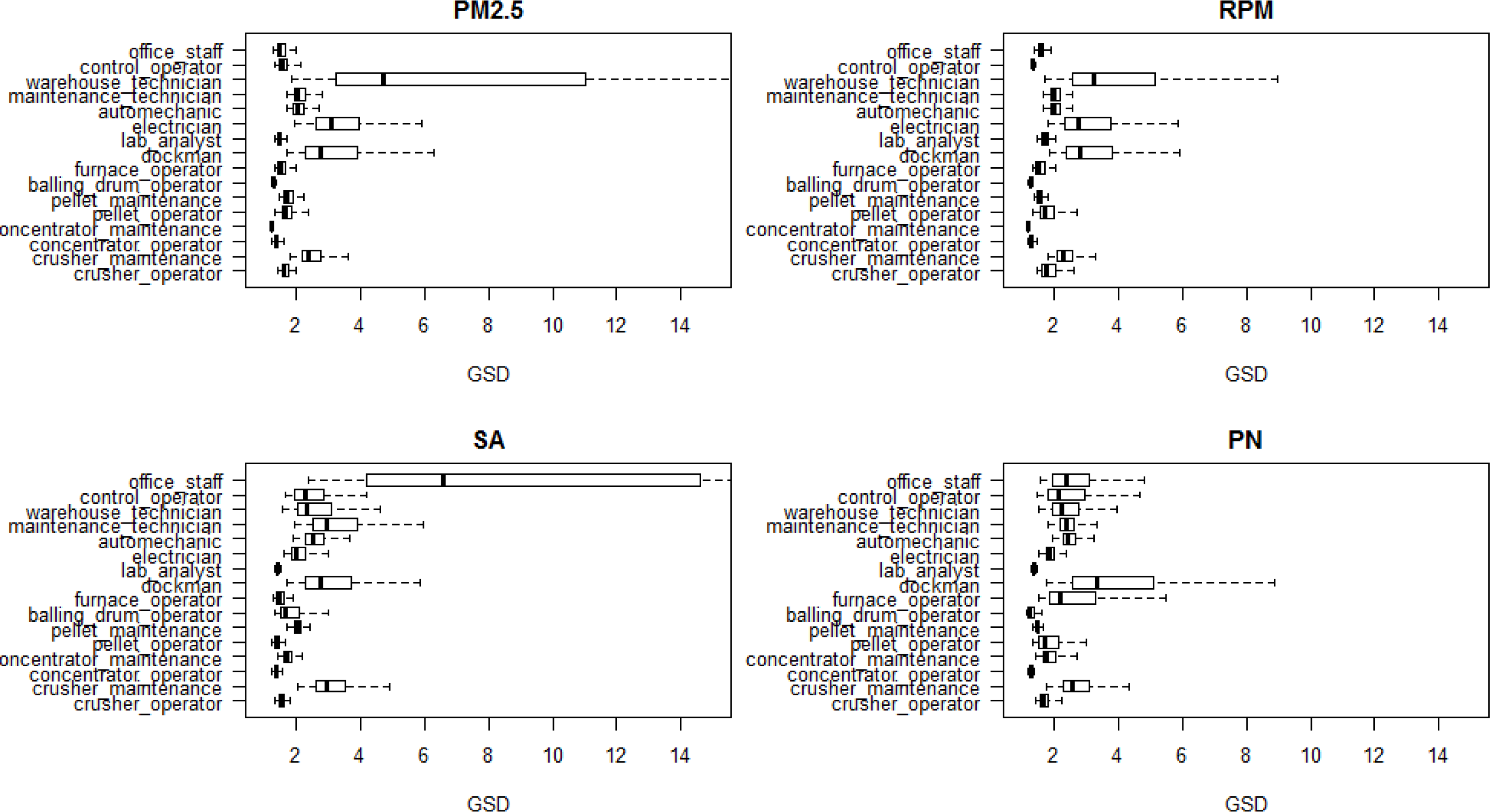
Boxplot of the geometric standard deviation within locations (GSD_WL) distributions by PM_2.5_, respirable particulate matter (RPM), alveolar-deposited surface area (ADSA), and particular number concentration and by exposure groups (SEG).

**Figure 5:**
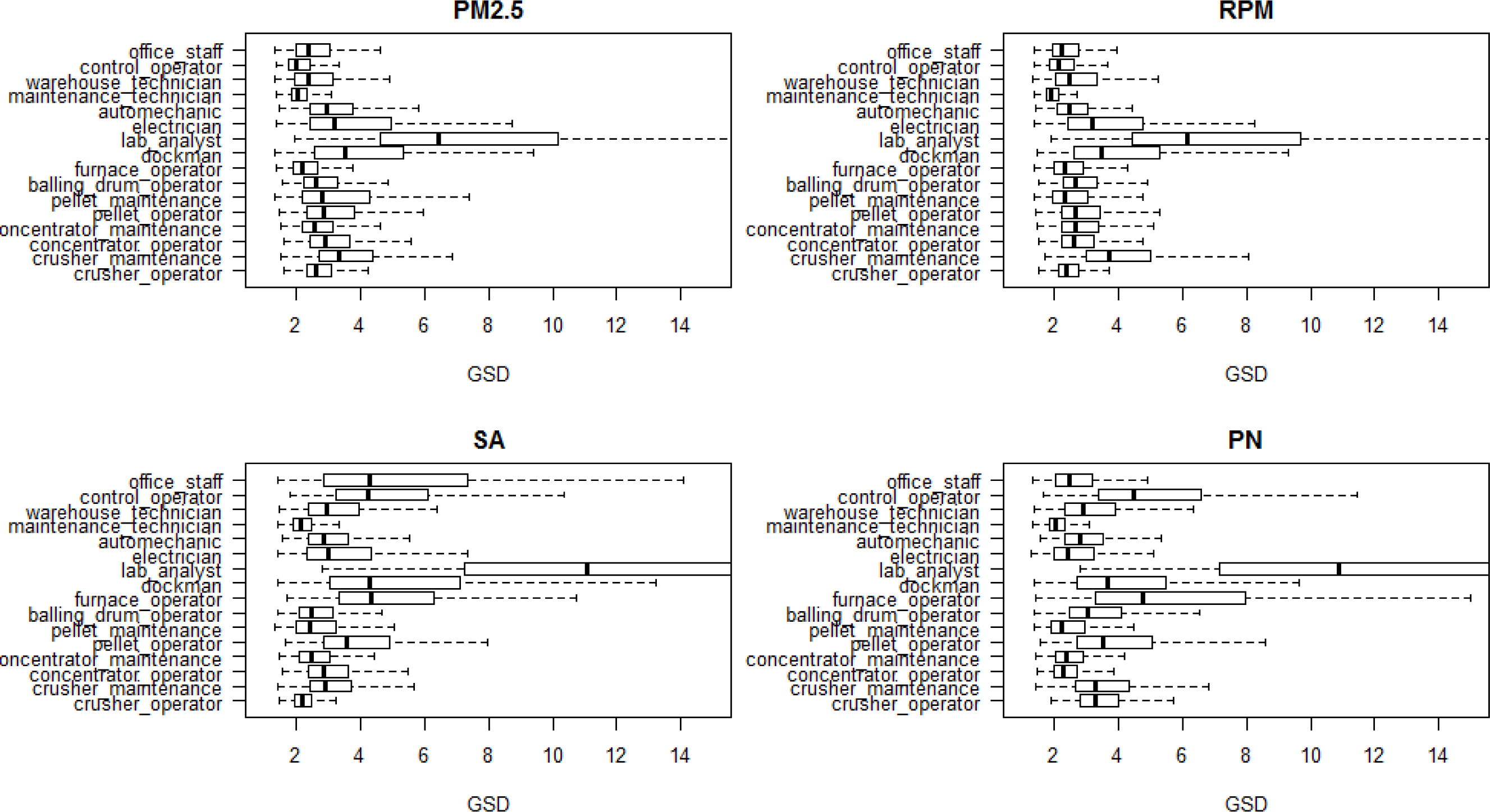
Boxplot of the geometric standard deviation between locations (GSD_BL) distributions by PM_2.5_, respirable particulate matter (RPM), alveolar-deposited surface area (ADSA), and particular number concentration and by exposure groups (SEG).

### Particle size distribution

Figure 6A shows that all SEGs had highest count fraction/µm in the smallest observed 0.02 – 0.3 μm size range. In contrast, the mass fraction/µm (converted from the size distribution count) was highest in the largest 5-10 µm bin (Figure 6B).

**Figure 6A:**
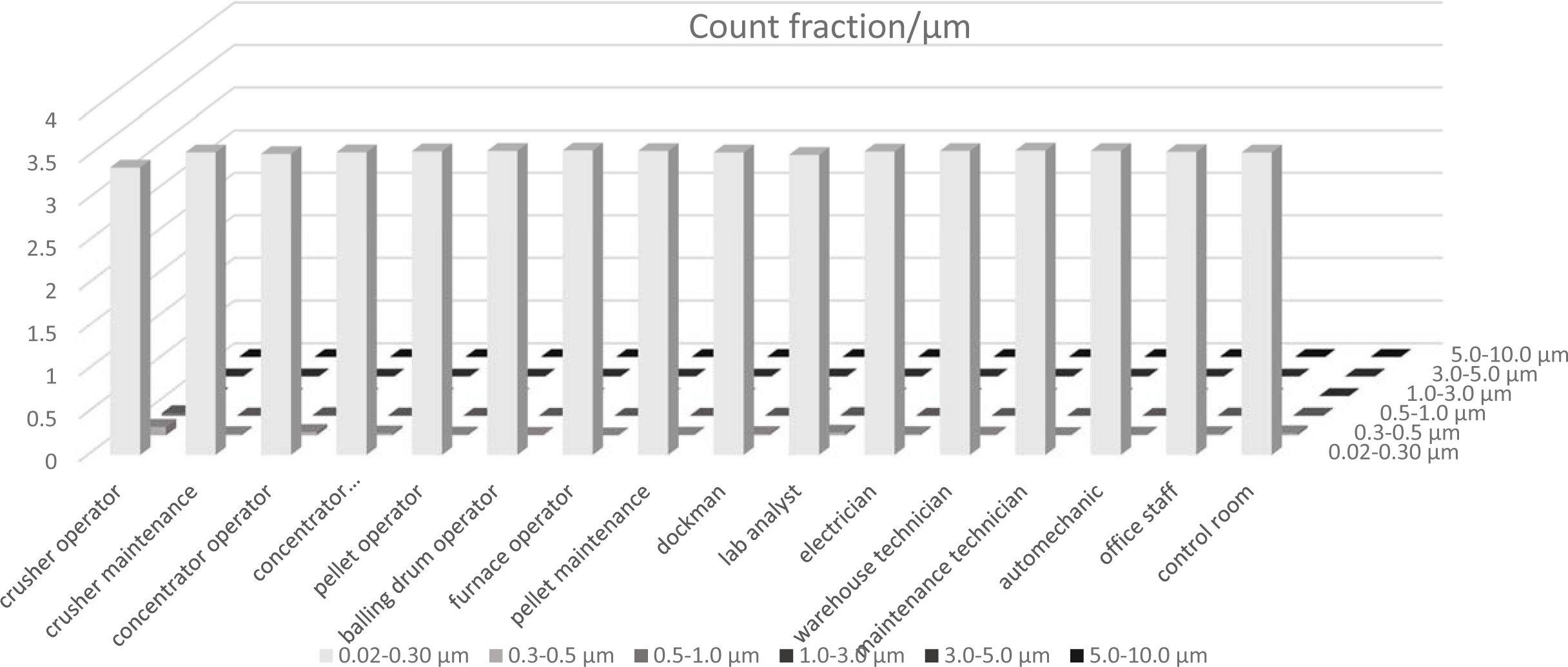
Distribution of the count fraction/µm by size bin and by SEG.

**Figure 6B:**
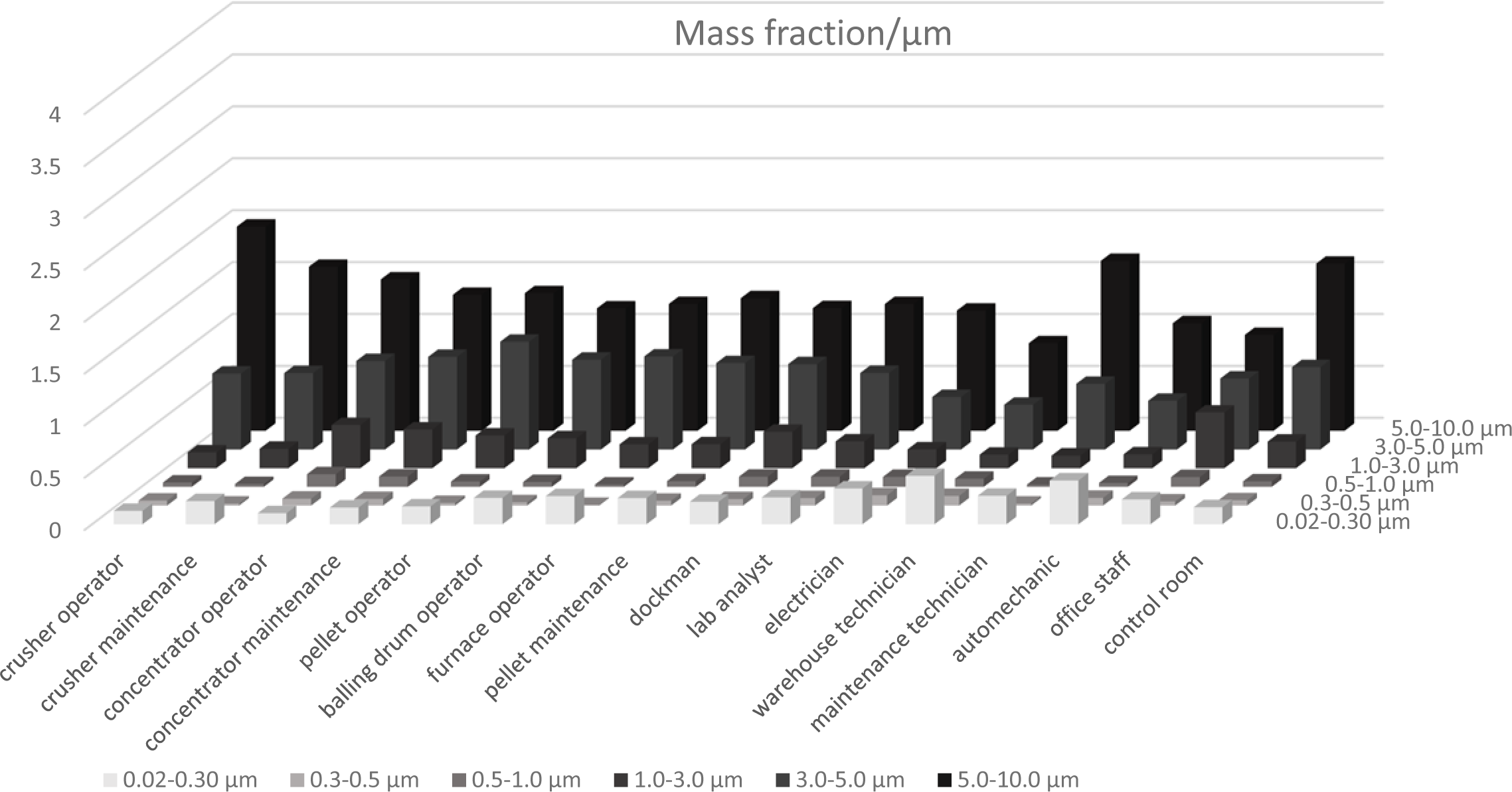
Distribution of the mass fraction/µm by size bin and by SEG.

## Discussion

### Exposure levels by SEG, mine, and metri**c**

Our study demonstrated the utility of direct-reading instruments to estimate unique characteristics of fine aerosol levels in various work areas across six taconite mines. The dust exposures at measured locations were lower than the ACGIH TLV for respirable dust (3 mg/m^3^). As expected, PM_2.5_ and RPM concentration tend to track each other. Highest RPM dust exposures were observed in areas within the pelletizing and concentrator departments (e,g, balling drum, pelletizing, furnace, concentrator operator), a trend that is generally consistent with the personal respirable dust exposures assessed by Hwang et al. (2017) in the same six mines. While the current study found *pelletizing maintenance* SEG had the highest exposure (0.38 mg/m^3^), the assessment with personal respirable samples found *balling drum operator* had the highest respirable dust exposure. The area RPM trend was not consistent with personal respirable silica trend found in Hwang et al. (2016) where the highest average personal respirable silica exposure was in the crusher department. The difference between the personal and the area monitor results could be due several reasons: 1) direct reading instruments cannot differentiate silica from other type of particles, 2) workers tend to move to various places within the department.

Very high SA and particle number concentrations were found in the concentrating and pelletizing processes where very high number of fine particles were released during the rolling and tumbling of the powder and pellets. These findings were consistent with the high correlation of SA and PA presented in Figure 2. Heitbrink et al (2006) performed a real-time monitoring study at an automotive engine manufacturing plant and learned that correlation for SA and PN were stronger during the winter when they used direct-fire gas fired heaters that generated substantial amount of ultrafine particles compared to the summer months. The strong correlation between SA and PA in this study suggested the presence of ultrafine particles in many locations in taconite operations. In some locations, we found high mass concentration ranking did not result in high surface area and particle number concentration ranking and vice versa (eg., concentrator operator, furnace operator) which aligns with our current theoretical knowledge that relatively high numbers of large particles do not contribute much to the overall surface area nor overall particle number. Some locations (e.g., pelletizing maintenance), however, may have both very large particles and ultrafine particles (e.g, balling drum operator has 4 metrics in top 3 ranking). Thus, it may be important to measure a wide range of particle sizes in order to better characterize the nature of the air pollution in these locations. PN does not correlate very well with RPM and PM_2.5_ (0.31 and 0.30, respectively) and the correlation between SA and RPM or PM_2.5_ were just slightly better (0.44, and 0.40, respectively). This finding suggests that SA and PN may be used interchangely to characterize ultrafine particles exposure, and PM_2.5_ or RPM may underestimate their presence. In certain scenarios such as nanomaterials monitoring where surfaces are known to be coated with different materials which might enhance the toxicity of the particles, surface area monitoring may be the better choice.

Despite the lack of motivation for employers to monitor fine particles for compliance purposes, our findings have important implications for epidemiological studies because the choice of metric can affect the ability to estimate disease risk and the characteristics of the exposure-response relationships. While we detected high mass concentrations in some areas within the pelletizing and concentrating departments, we also found high surface area and particle number concentrations which indicated the presence of very fine particles concentrated in the pelletizing department. Thus, the use of mass metric may potentially attenuate the dose-response for cardiovascular risk from fine particle exposure in some settings.

### Within and between-location variability by SEG, mine, and metric

For all metrics, we observed lower within-location variability (GSD_WL) in the crusher (except crusher maintenance), concentrator, and pelletizing departments (except dockman) where many large taconite processing machines (crushers, rod mills, ball mills, etc..) ran continuously throughout the day compared to the shop areas where intermittent maintenance activities usually took place. For example, forklifts in the warehouse generated large numbers of fine particles; as did moving the trucks in and out of the shop and welding activities in the truck shops. In most cases (except the *crusher operator* and the *maintenance technician),* we only sampled one location at each mine for each SEG so the between-location variability is essentially the between-mine variability. Large between-location variability (GSD_BL) indicates mine-specific differences in the taconite mining process and it could be due to a number of factors including size of each mine (1400 workers vs less than 100), rock formation that changes from soft to hard, and slight differences in the job descriptions (Hwang et al., 2016). Other factors that might contribute to the observed mine-specific differences could be unique events during the sampling day (e.g., a certain process being shut down for maintenance). For example, there was nobody working in one of the labs during a sampling period. We also noticed that the GSD_BL and their uncertainties tend to be higher in some relatively clean areas such as the office, control room, lab, shop areas, pellet docking area for SA and PN compared to PM_2.5_ and RPM. It could be that while these locations look relatively less dusty, the SA and PN monitors can pick up more fine particles than the PM_2.5_ and RPM.

### Particle size distribution

Because most of fine particles were in the 0.02-0.3 range across all SEGs, it was difficult to distinguish between the SEGs. There was more variability between the bins and between groups in terms of mass concentrations. The size distribution analysis corroborated our particle number and surface area monitoring data by demonstrating the presence of ultrafine particle numbers in taconite mining operations. While our current data did not have more refined particle size distribution within the 0.02-0.3 µm range, understanding of the particle size distribution is important in epidemiology studies because it would enable us to identify which particle sizes may be more relevant to toxicity than others. For example, Oberdorster et al. (1994) found that titanium dioxide particles at around 20 nm cleared significantly slower and translocated to more interstitial sites and regional lymph nodes compared to ∼200 nm particles in rats.

### Statistical analysis of real-time data

A common statistical method for analyzing real-time exposure data is some form of autoregressive model to account for temporal correlation (e.g, Entink et al., 2011). That being said, the assumptions underlying autoregressive processes may be violated in the presence of factors that result in sudden spikes or dips in concentrations. The effects of these spikes/dips can be mitigated if these factors are known and included as covariates in the statistical model. Another approach to analyzing real-time data is to evaluate peak exposures but peak definition has been somewhat arbitrary depending on the study. While Preller et al. (2004) defined peak exposure as when the concentration exceeded the time weighted average over the monitoring period (reference level), Meijster et al. 2007 chose a reference level that is approximately equal to the population average exposure in the flour processing industry. Other studies (e.g., Nieuwenhuijsen et al., 1995) defined peak exposure in their study of respiratory health effects among bakers as the highest exposure measured during a specific task within a group of workers. While evaluating peak exposure is beyond the scope of this study, we recognize the importance of peak exposures and its influence on the statistical models.

*Limitations:* Our monitoring campaign captured a snap shot of particle exposures at various locations where our SEGs typically work; thus it is not representative of the personal exposures. While some groups of workers will spend more time at their defined locations (e.g., office workers, control room operators, auto mechanics), other groups move around to various parts of the buildings (e.g., crusher operators, crusher maintenance) which would make the personal exposures very different from area dust concentrations. Furthermore, in most cases, due to time constraints, we were able to sample only one location per SEG and only in the morning at each mine. This limited our ability to characterize the variability for the entire 8-hour shift, day to day, and seasonal variations. As mentioned previously, the area of each workplace in some departments – such as the crusher, the concentrator, and pelletizing area – is enormously large compared to other places in the mine. One sample within that workplace might not adequately capture the spatio-temporal variability of exposure within one location and between locations within an SEG. It is also worth noting that when we ran the real-time monitors, we also ran a high-volume sampling pump for the NanoMOUDI impactors to collect elongate mineral particles on the filters at the same time and location. At a few small and relatively clean locations such as the office, dockman, and lab, we occasionally noticed some build-up of smoke and smell of oil at the end of the sampling period that came from the high volume sampling pump. While we think that close proximity to the pump might have affected the concentration at those locations, measurements at most other locations were unlikely to be affected substantially since neither the smell nor smoke was apparent to us.

Despite these limitations, the strength of our study is the large number of sampling locations and measurements of particle concentrations using four different metrics. These measurements allowed us to compare the concentrations across metrics and how the metrics influence the ranking of various job groups within the taconite mines. This information could potentially be used in epidemiological studies to explore the effects of fine aerosol exposures on cardiovascular health risks among taconite miners.

## CONCLUSION

Our study used an array of real-time instruments to characterize fine aerosol concentrations using multiple exposure metrics at various locations across six taconite mines in Northern Minnesota. Our monitoring campaign identified locations with high particle exposures and showed that the rank of those high exposure groups change by metric. Typically the strategy for classifying similarly exposure groups might be influenced by the metric used, which would subsequently affect the validity of the epidemiological studies that used these exposure data. In this case, the classification of high exposure groups slightly changed based upon the mass, ADSA, or particle number metric. Particle number and surface area exposure metrics were highly correlated with each other. This study highlights the importance of considering alternative exposure metrics such as alveolar-deposited surface area and particle number concentration along with mass concentration to study the health effects from fine aerosol exposures in occupational epidemiology studies at taconite mines.

## ACKNOWLEDGMENT

This study was funded by the State of Minnesota. The conclusions belong to the authors and do not reflect the State of Minnesota. The authors would also like to thank Rob Kelly for their assistance in the data collection as well as the taconite mining companies and employees for their help in collecting the data.

## References

Allen EM, Alexander BH, MacLehose RF, Ramachandran G, Mandel J. (2014) Mortality experience among Minnesota taconite mining industry workers. Occupational & Environmental Medicine, 71(11), pp.744–749.

Allen EM, Alexander BH, MacLehose RF, Nelson HH, Ramachandran G, Mandel JH. (2015) Cancer incidence among Minnesota taconite mining industry workers. Ann Epidemiol. Nov;25(11):811–5. doi: 10.1016/j.annepidem.2015.08.003.

Allen EM, Alexander BH, MacLehose RF, Nelson HH, Ryan AD, Ramachandran G, Mandel JH. (2015) Occupational exposures and lung cancer risk among Minnesota taconite mining workers. Occup Environ Med. Sep;72(9):633–9.doi: 10.1136/oemed-2015-102825.

Astrakianakis G, Seixas N, Camp J et al (2006), Modeling, estimation, and validation of cotton dust and endotoxin exposures in Chinese textile operations. Ann Occup Hyg: Ann Occup Hyg 2006; 50(6): 573–582.doi: 10.1093/annhyg/mel018

Axten C and Foster D (2008) Analysis of airborne and waterborne particles around a taconite ore processing facility. Re Regul.Toxicol. Pharmacol. 52(1 Suppl):S66–72.

Baldauf, RW., Devlin, R. B., Gehr, P., Giannelli, R., Hassett-Sipple, B., Jung, H., Walker, K. (2016). Ultrafine Particle Metrics and Research Considerations: Review of the 2015 UFP Workshop. International Journal of Environmental Research and Public Health, 13(11), 1054. http://doi.org/10.3390/ijerph13111054

Berndt, ME, and Brice WC (2008) The origins of public concern with taconite and human health: Reserve mining and the asbestos case. Regul.Toxicol. Pharmacol. 52(1 Suppl):S31–S39(2008).

Carlin BP, Louis TA. (2008) Bayesian methods for data analysis. 3rd edn. Boca Raton, FL: Chapman & Hall/CRC.

Case, BW, Abraham JL, Meeker G, Pooley FD, Pinkerton KE (2011) Applying definitions of “asbestos” to environmental and “low dose” exposure levels and health effects, particularly malignant mesothelioma. J. Toxicol. Environ. Health, Part B 14:3–39

Chung A, Chang DPY, Kleeman MJ, Perry KD, Cahill TA, Dutcher D, McDougall EM, Stroud K (2001) Comparison of real-time instruments used to monitor airborne particulate matter Journal of the Air and Waste Management Association, 51 (2001), pp. 109–120

Clark, T, Harrington, V, Morgan, AJ, Sargent, E (1980). Respiratory effects of exposure to dust in taconite mining and processing. Am Rev Respir Dis., 121(6), 959–966.

Cooper WC, Wong O, Graebner R. (1988) Mortality of workers in two Minnesota taconite mining and milling operations. J Occup Med. 30:506–11

Cooper WC, Wong O, Trent LS, et al..(1992) An updated study of taconite miners and millers exposed to silica and non-asbestiform amphiboles. J Occup Med. 34:1173–80.

Costello S., Daniel M Brown, Elizabeth M Noth, Linda Cantley, Martin D Slade, Baylah Tessier-Sherman, S Katharine Hammond, Ellen A Eisen, and Mark R Cullen, “Incident ischemic heart disease and recent occupational exposure to particulate matter in an aluminum cohort,” Journal of Exposure Science and Environmental Epidemiology, no. 1. pp. 82–88(2014).

Dockery DW, Pope CA, Xu X, et al.. An Association between Air Pollution and Mortality in Six U.S. Cities. N Engl J Med. 329:1753–9(1993).

Dockery, D., Luttmann-Gibson, H., Rich, D., Link, M., Mittleman, M., Gold, D., Verrier, R. (2005). Association of Air Pollution with Increased Incidence of Ventricular Tachyarrhythmias Recorded by Implanted Cardioverter Defibrillators. Environmental Health Perspectives, 113(6), 670–674. Retrieved from http://www.jstor.org/stable/3436292

Entink, RK, Fransman W, and Brouwer D (2011) How to statistically analyze nano exposure measurement results: Using an ARIMA time series approach. Journal of Nanoparticle Research 12(12)DOI: 10.1007/s11051-011-0610-x

Environmental Protection Agency. Particulate Matter. Available at http://www.epa.gov/air/particlepollution/health.html (Accessed May 7, 2017).

Environmental Protection Agency. Taconite Iron Ore NESHAP Economic Impact report. 2003. Available at http://www.epa.gov/ttn/atw/taconite/taconite_eia.pdf (Accessed May 7, 2017)

Gallagher, LG, Ray, RM, Li, W, Psaty, BM, Gao, DL, Thomas, DB and Checkoway, H. (2012), Occupational exposures and mortality from cardiovascular disease among women textile workers in Shanghai, China. Am. J. Ind. Med., 55: 991–999.doi:10.1002/ajim.22113

Gamble, J. F., and G. W. Gibbs: An evaluation of the risks of lung cancer and mesothelioma from exposure to amphibole cleavage fragments. Regul. Toxicol. Pharmacol. 52(1 Suppl):S154–S186(2008).

Heitbrink WA, Evans DE, Ku BK, Maynard AD, Slavin TJ, Peters TM. (2009)Relationships Among Particle Number, Surface Area, and Respirable Mass Concentrations in Automotive Engine Manufacturing. J Occup Environ Hyg. 2009 Jan;6(1):19–31.doi: 10.1080/15459620802530096.

Higgins, IT, Glassman JH, Oh MS et al (1983) Mortality of Reserve Mining Company http://www.dnr.state.mn.us/education/geology/digging/history.html. (Accessed May 6, 2017)

Hwang, J., Ramachandran, G., Raynor, P. C., Alexander, B. H., & Mandel, J. H. (2013). Comprehensive assessment of exposures to elongate mineral particles in the taconite mining industry. The Annals of Occupational Hygiene, 57(8), 966–78. http://doi.org/10.1093/annhyg/met026

Hwang, J, Ramachandran G, Raynor PC, Alexander BH, Mandel JH (2014). The relationship between various exposure metrics for elongate mineral particles (EMP) in the taconite mining and processing industry. Journal of occupational and environmental hygiene, 11(9), pp.613–24. Available at: http://www.ncbi.nlm.nih.gov/pubmed/24512074.

Hwang, J.,(2016). Assessing present-day and historical exposures of workers to taconite dust in the iron mining industry in Northeastern Minnesota. Doctoral dissertation. University of Minnesota.https://conservancy.umn.edu/bitstream/handle/11299/158414/Hwang_umn_0130E_14041.pdf?sequence=(Accessed May 6, 2017)

Lambert CS, Alexander BH, Ramachandran G, MacLehose RF, Neslon HH, Ryan AD, Mandel JH. 2015. A case-control study of mesothelioma in Minnesota iron ore (taconite) miners. Occupational and environmental medicine, 73(2), pp.1–7. Available at: http://oem.bmj.com/lookup/doi/10.1136/oemed-2015-103105

Mandel, J.H., Ramachandran, G. & Alexander, B.H., 2016. Increased Lung Cancer Mortality in Taconite Mining: The Potential for Disease from Elongate Mineral Particle Exposure. Chemical Research in Toxicology, 29(2), pp.136–141. Available at: http://pubs.acs.org/doi/abs/10.1021/acs.chemrestox.5b00472 (Accessed May 7, 2017)

Plummer M (2003). JAGS: A Program for Analysis of Bayesian Graphical Models Using Gibbs Sampling, Proceedings of the 3rd International Workshop on Distributed Statistical Computing (DSC 2003), March 20–22, Vienna, Austria. ISSN 1609-395X

Maynard, AD, Maynard RL(2002) A derived association between ambient aerosol surface area and excess mortality using historic time series data. Atmospheric Environem. 36, 5561–5567.

Maynard AD, Aitken RJ (2007). Assessing exposure to airborne nanomaterials. Current ability and future requirements. Nanotoxicology. 1.26–41.

Meier R, Cascio WE, Ghio AJ, Wild P, Danuser B, Riediker M. 2014. Associations of short-term particle and noise exposures with markers of cardiovascular and respiratory health among highway maintenance workers. Environ Health Perspect 122:726–732; http://dx.doi.org/10.1289/ehp.1307100

Methner M, Beaucham C, Crawford C, Hodson L, & Geraci C. (2012). Field Application of the Nanoparticle Emission Assessment Technique (NEAT): Task-Based Air Monitoring During the Processing of Engineered Nanomaterials (ENM) at Four Facilities. JOEH. 9(9):543–555.doi:10.1080/15459624.2012.699388

Mine Safety and Health Administration (MSHA). Mine Data Retrieval System. https://arlweb.msha.gov/drs/drshome.htm

Minnesota Department of Health (1999) Cancer incidence rates in Northeastern Minnesota. In Minnesota Cancer Surveillance System Epidemiology Report. http://www.health.state.mn.us/divs/hpcd/cdee/mcss/documents/necancer99-2.pdf (Accessed May 6, 2017)

Minnesota Department of Health (2003). Cancer incidence rates in Northeastern Minnesota with an emphasis on mesothelioma. In Minnesota Cancer Surveillance System Epidemiology Report. http://www.health.state.mn.us/divs/hpcd/cdee/mcss/documents/NEMNCancerMeothelioma031.pdf (Accessed May 6, 2017)

Minnesota Department of Natural Resources. Minnesota mining history. Available at http://www.dnr.state.mn.us/education/geology/digging/history.html (Accessed May 7, 2017)

Meijstern T, Tielemans E, Chinkel J, and Heederik D (2008) Evaluation of peak exposures in the Dutch flour processing industry: implications for intervention strageties. Ann. Occup. Hyg., Vol 52, 7, p587–596.

Neophytou AM, Costello S, Brown DM, Picciotto S, Noth EM, Hammond SK, Cullen MR, Eisen EA; Marginal Structural Models in Occupational Epidemiology: Application in a Study of Ischemic Heart Disease Incidence and PM2.5 in the US Aluminum Industry. Am J Epidemiol 2014; 180(6): 608–615.doi: 10.1093/aje/kwu175

Neophytou AM, Noth EM, Liu S, Costello S, Hammond SK, Cullen MR, et al.. (2016) Ischemic Heart Disease Incidence in Relation to Fine versus Total Particulate Matter Exposure in a U.S. Aluminum Industry Cohort. PLoS ONE 11(6): e0156613.doi:10.1371/journal.pone.0156613

Nieuwenhuijsen MJ, Sandiford CP, Lowson D et al. (1995) Peak exposure concentrations of dust and flour aeroallergen in flour mills and bakeries. Ann Occup Hyg; 39:193–201.

Oberdörster, G., Oberdörster, E., & Oberdörster, J. (2005). Nanotoxicology: An Emerging Discipline Evolving from Studies of Ultrafine Particles. Environmental Health Perspectives, 113(7), 823–839. http://doi.org/10.1289/ehp.7339

Oberdorster G, Ferin J, Lehnert BE. Correlation between particle size, in vivo particle persistence, and lung injury. Environ Health Perspect. 1994;102:173–179. [PMC free article] [PubMed]

Odo, NU, Mandel J, Perlman, DM, Alexander BH, Scanlon PD. (2013). Estimates of restrictive ventilatory defect in the mining industry. Considerations for epidemiological investigations: a cross-sectional study. BMJ open, 3(7).

Park J.Y. P.C. Raynor, G. O. Eberly, G. Ramachandran (2010) Comparing exposure groups by different exposure metrics using statical parameters:contrast and precision. Ann Occup Hyg 54(7):799–812

Pope CA III, Burnett RT, Thurston GD, et al.. Cardiovascular mortality and long-term exposure to particulate air pollution: epidemiological evidence of general pathophysiological pathways of disease. Circulation. 109:71–7(2004).

Preller L, Burstyn I, de Pater N et al. (2004) Characteristics of peaks of inhalation exposure to organic solvents. Ann Occup Hyg; 48: 643–52.

R Core Team (2016). R: A language and environment for statistical computing. R Foundation for Statistical Computing, Vienna, Austria. URL https://www.R-project.org/.

Ramachandran G, Paulsen, D, Watts, W., Kittelson, D (2005). Mass surface area and number metrics in diesel occupational exposure assessment. J. Envrion. Monit, 7, 728–735. DOI: 10.1039/b503854e

Reto Meier, Wayne E. Cascio, Brigitta Danuser, Michael Riediker; Exposure of Highway Maintenance Workers to Fine Particulate Matter and Noise. Ann Occup Hyg 2013; 57(8): 992–1004.doi: 10.1093/annhyg/met018

Schwartz J, Marcus A. (1990) Mortality and air pollution in London: Atime series analysis.AmJ Epidemiol; 131: 185–94

Sheehy J. Reconstruction of occupational exposures to silica containing dusts in the taconite industry. Doctoral dissertation. Minneapolis, MN: University of Minnesota. (1986)

Toren K, I.A., Bergdahl, T. Nilsson, B. Jarvholm. Occupational exposure to particulate air pollution and mortality due to ischaemic heart disease and cerebrovascular disease. Occup Environ Med. Occup Environ Med 2007;64:515–519.

University of Minnesota, 2014. Public Meeting and Final Report December 1, 2014. Available at: http://taconiteworkers.umn.edu/news/pages/dec12014_mtg.html [Accessed May 6, 2017].

Wilson R, McConnell EE, Ross M et al. (2008) Risk assessment due to environmental exposures to fibrous particulates associated with taconite ore. Regul Toxicol Pharmacol; 52 (1 Suppl.): S232–45.

Yanoksy, J.D., Williams, P.L., and MacIntosh, D.L. 2002. A Comparison and Two Direct-Reading Aerosol Monitors with the Federal Reference Method for PM2.5 in Indoor Air. Atmos. Environm. 36.107–113.

